# Extremely fast and accurate open modification spectral library searching of high-resolution mass spectra using feature hashing and graphics processing units

**DOI:** 10.1101/627497

**Authors:** Wout Bittremieux, Kris Laukens, William Stafford Noble

## Abstract

Open modification searching (OMS) is a powerful search strategy to identify peptides with any type of modification. OMS works by using a very wide precursor mass window to allow modified spectra to match against their unmodified variants, after which the modification types can be inferred from the corresponding precursor mass differences. A disadvantage of this strategy, however, is the large computational cost, because each query spectrum has to be compared against a multitude of candidate peptides.

We have previously introduced the ANN-SoLo tool for fast and accurate open spectral library searching. ANN-SoLo uses approximate nearest neighbor indexing to speed up OMS by selecting only a limited number of the most relevant library spectra to compare to an unknown query spectrum. Here we demonstrate how this candidate selection procedure can be further optimized using graphics processing units. Additionally, we introduce a feature hashing scheme to convert high-resolution spectra to low-dimensional vectors. Based on these algorithmic advances, along with low-level code optimizations, the new version of ANN-SoLo is up to an order of magnitude faster than its initial version. This makes it possible to efficiently perform open searches on a large scale to gain a deeper understanding about the protein modification landscape. We demonstrate the computational efficiency and identification performance of ANN-SoLo based on a large data set of the draft human proteome.

ANN-SoLo is implemented in Python and C^**++**^. It is freely available under the Apache 2.0 license at https://github.com/bittremieux/ANN-SoLo.

## 1 Introduction

As mass spectrometry (MS) instrumentation has matured over the last decade, the focus of proteomics experiments has shifted from identifying the peptides and proteins that are present in a biological sample to characterizing all proteoforms therein [1]. Detection of proteoforms yields additional biological insights relative to simple peptide or protein lists, because the proteoforms capture disparate sources of biological variation that alter the primary protein sequences, such as post-translational modifications (PTMs) and amino acid mutations.

Accordingly, detecting proteoforms in MS requires detecting PTMs, which can be challenging. In particular, appropriate search settings are needed to correctly identify spectra corresponding to modified peptides. A common approach is to specify the variable modifications that are expected to be present a priori. A downside of this approach, however, is that as the number of potential modifications increases the search space explodes, leading to long search times and reduced identification sensitivity. A compounding problem is that, besides modifications of biological interest, other modifications can be introduced during the various sample processing steps as well [2]. This can lead to challenges to untangle these artificial modifications from the interesting modifications.

An alternative to explicitly specifying a limited number of variable modifications is open modification searching (OMS) [3, 4]. OMS works by using a very wide precursor mass window, exceeding the delta mass induced by PTMs, to infer identifications of modified spectra from partial matches against their unmodified variants. Afterwards, the presence and types of the modifications can be inferred from the differences between the observed precursor masses and the masses of the unmodified peptides [5]. In this fashion, all possible modifications are implicitly considered, allowing an untargeted analysis of all modifications that are present.

A downside of using a very wide precursor mass window, however, is that the search space is considerably enlarged relative to a standard database search, rendering OMS computationally expensive. As a result, historically OMS has only been used to a limited extent and with severely restricted protein databases. Based on computational and algorithmic advances, however, recently several modern open search engines have been developed that can efficiently handle this large search space [6–12]. These tools make it possible to perform OMS on a proteome-wide scale, allowing researchers to gain a deeper understanding of the protein modification landscape.

Here we present an update to our **A**pproximate **N**earest **N**eighbor **S**pectral **L**ibrary (ANN-SoLo) tool for efficient open modification spectral library searching [8]. As described previously, ANN-SoLo uses approximate nearest neighbor (ANN) indexing to speed up OMS by selecting only a limited number of the most relevant library spectra to compare to an unknown query spectrum. This approach is combined with a cascade search strategy [13] to maximize the number of identified unmodified and modified spectra while strictly controlling the false discovery rate (FDR). Additionally, the shifted dot product score is used to sensitively match modified spectra to their unmodified counterparts by taking both directly matching fragments and fragments that match according to the precursor mass difference into account [8].

We describe two major improvements to ANN-SoLo. First, we show how feature hashing is used to convert high-resolution tandem mass spectrometry (MS/MS) spectra to vectors with a limited dimensionality while closely capturing the high fragment resolution. Hashed spectrum vectors approximate the original spectra better compared to simply binning the spectra to vectors, leading to an improvement in accuracy of the ANN candidate selection step. Second, the spectral library candidate selection step is sped up by using specialized graphics processing unit (GPU) hardware. Whereas GPUs have previously been proposed to accelerate spectral matching [14–16], to our knowledge this work is the first application of GPUs to efficiently process large search spaces, such as during OMS. We show how these two developments increase the speed of ANN-SoLo by up to an order of magnitude, making it possible to perform OMS extremely efficiently. We demonstrate this high computational performance through an open search of a large data set of the draft human proteome and investigate human PTMs.

ANN-SoLo is implemented in Python and C^**++**^. It is freely available as open source under the permissive Apache 2.0 license at https://github.com/bittremieux/ANN-SoLo.

## 2 Methods

### 2.1 Feature hashing to vectorize high-resolution mass spectra

To build an ANN index to efficiently select candidates from the spectral library, spectra are vectorized to represent them as points in a multidimensional space. In previous work [8], we converted spectra to sparse vectors by dividing the mass range into equally spaced bins and assigning each peak’s intensity to the corresponding bin. When choosing the mass bin width two conflicting factors must be considered. First, the mass bins should be as small as possible, ideally corresponding to the fragment mass tolerance, to accurately capture the peak masses. Second, because the sensitivity of multidimensional indexing techniques decreases as the dimensionality increases, due to the curse of dimensionality [17], shorter vectors are preferred. Previously, we empirically found that mass bins of 1 Da represented a good trade-off between fragment mass resolution and vector dimensionality [8]. However, because such mass bins considerably exceed the fragment mass tolerance when dealing with high-resolution spectra, multiple distinct fragments occasionally get merged into the same mass bin. This merging leads to an overestimation of the spectral similarity when comparing two spectra with each other using their vector representations, because spurious matches between fragments can occur.

In this work, we employ a different vectorization scheme. Rather than binning spectra to vectors directly, we use feature hashing [18] to convert high-resolution spectra to low-dimensional vectors (figure 1). The following two-step procedure is used to convert a high-resolution MS/MS spectrum to a vector:

**Figure 1:**
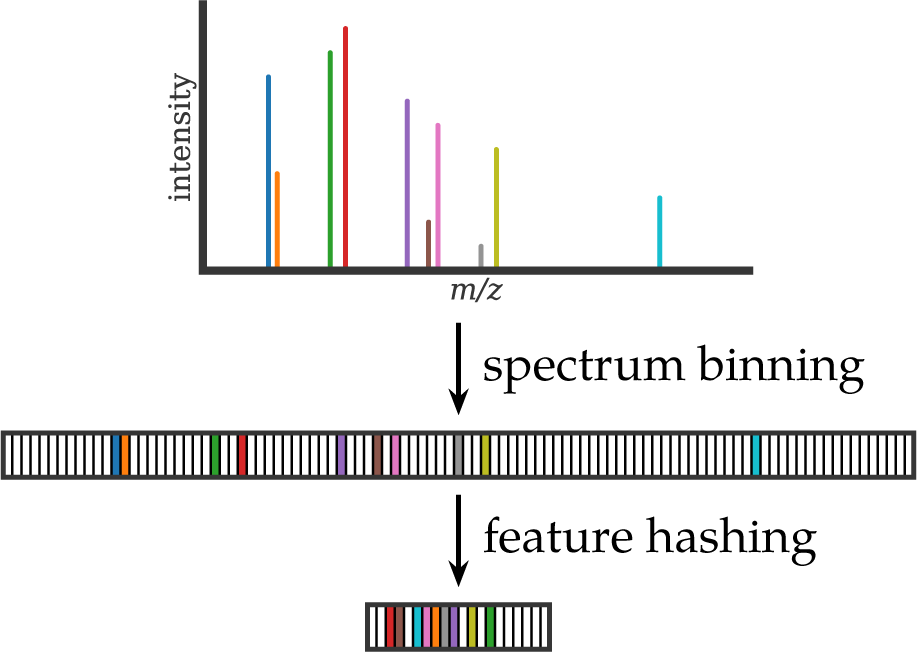
High-resolution MS/MS spectra are first converted to sparse vectors using small mass bins to accurately capture the fragment masses. Next, these high-dimensional, sparse vectors are converted to lower-dimensional vectors through feature hashing.

1. Convert the spectrum to a sparse vector using small mass bins to tightly capture fragment masses.
2. Hash the sparse, high-dimensional vector to a lower-dimensional vector by using a hash function to map the mass bins to a limited number of hash bins.

More precisely, let *h*: ℕ → {1,…, *m*} be a random hash function. Then *h* can be used to convert a vector *x* = ⟨*x*_1_,…, *x*_*n*_⟩ to a vector 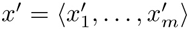, with *m* ≪ *n*:

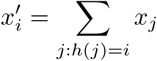

It can be proven that under moderate assumptions feature hashing approximately conserves the Euclidean norm [19], and hence, the similarity between hashed vectors can be used to approximate the similarity between the original, high-dimensional vectors.

An important consideration in choosing hash function *h* is that it must be unbiased in order to minimize the number of hash collisions. Hash collisions occur when fragment peaks in multiple distinct mass bins are mapped to the same hash bin. This can happen because the number of hash bins is significantly lower than the number of original mass bins. Hash collisions cause unrelated fragment peaks to be matched with each other, leading to an inflated similarity between two spectrum vectors. To avoid a systematic hash collision bias hash function *h* has to be truly random. This is, for example, not the case for the classic multiply-mod-prime hashing scheme [20]. In this work, we instead use the MurmurHash3 algorithm [21], a popular non-cryptographic hash function that essentially behaves as truly random hashing [20]. MurmurHash3 is an efficient, general-purpose hash function that uses multiplications, rotations, XOR operations, and bit shifts to convert an input key to a random hash value. Subsequently, the modulo operator is used to restrict the hash value to a user-specified number of hash bins. The number of hash bins *m*, i.e. the length of the hashed vectors, directly influences the rate of collisions. The smaller *m* is, the more likely it is that two or more fragment peaks will be mapped to the same hash bin. Nevertheless, for a suitable value of *m*, because spectra contain only a small number of fragment peaks, the unhashed spectrum vectors are very sparse and can be converted to low-dimensional vectors without suffering too many hash collisions.

### 2.2 GPU-powered spectral library candidate selection

ANN-SoLo uses approximate nearest neighbor indexing to efficiently find the relevant candidate spectra in the spectral library that need to be matched against each query spectrum when performing an open search [8]. First, an ANN index is constructed using the vectorized library spectra. Subsequently, to identify an unknown query spectrum, a nearest neighbor search using the ANN index is performed to select a limited set of library candidates. Finally, the optimal match between the query spectrum and its library candidates is computed using the shifted dot product to accurately match modified spectra to their unmodified library counterparts.

Typically, during an open search each query spectrum has to be compared to most of the spectra in the library, imposing a significant computational burden. In contrast, ANN-SoLo massively speeds up open searches by using an ANN index to efficiently select only a limited number of relevant library candidates to be evaluated for each query spectrum. Whereas previously the Annoy library [22] was used for ANN searching, this is now done using the Faiss library [23], developed by Facebook for web-scale similarity searching. A major advantage of Faiss is that it can use NVIDIA CUDA-enabled GPUs to accelerate ANN searching, resulting in significant speed-ups [24].

ANN searching in Faiss is based on the concept of an inverted index [25]. To construct the ANN index a representative set of vectors is selected by using the centroids of a k-means clustering operation. Next, for each of these centroids a list of references to the library spectra that are closest to it is stored in an inverted index file. Subsequently, during ANN querying, finding the candidates for a query spectrum no longer requires searching the entire spectral library. Instead, the query only needs to be compared against the small number of centroids in the inverted index to retrieve the closest library candidates. The accuracy and speed of ANN indexing is governed by two hyperparameters: the number of lists used to partition the spectral library during index construction and the number of lists to probe during querying. Using a higher number of lists results in a more fine-grained partitioning of the data space, whereas probing more lists during querying decreases the chance of missing the best library candidate at the expense of running time.

### 2.3 Miscellaneous improvements

In addition to the important algorithmic changes described above, we have implemented several additional, smaller improvements.

#### Spectral library reading

A significant part of the ANN-SoLo runtime consists of reading spectra from the spectral library file. Because library spectra are retrieved from disk in an indeterminate order, based on the query spectra that are being identified, a large number of random-access read operations are needed. To optimize spectral library reading Cython [26] is used to parse the spectral library file efficiently by avoiding overhead from the Python IO libraries. Additionally, C-style memory mapping is used to perform random-access reads from binary spectral library files.

#### Spectrum preprocessing

To optimize the spectrum–spectrum match (SSM) scoring the query and library spectra are preprocessed to increase their signal-to-noise ratio. Spectrum preprocessing includes, for example, precursor peak and low-intensity noise peak removal, and fragment intensity scaling to de-emphasize overly dominant peaks. Although the previous spectrum preprocessing functionality was already implemented quite efficiently by making extensive use of the NumPy scientific Python library [27], it has been further optimized using Numba [28], a just-in-time compiler for Python. This preprocessing functionality has been extracted into the spectrum_utils software package [29] for general public use to preprocess and visualize MS/MS spectra.

#### Batch query processing

Rather than identifying each query spectrum individually and retrieving the candidate spectra from the ANN index for each query separately, the query spectra are now processed in batches. This makes it possible to exploit parallelism while querying the ANN index, optimally utilizing the GPU hardware to achieve a considerable speed-up.

### 2.4 Data sets

The main data set used to evaluate the improvements made to ANN-SoLo was generated in the context of the 2012 study by the Proteome Informatics Research Group of the Association of Biomolecular Resource Facilities, whose goal was to assess the community’s ability to analyze modified peptides [30]. The biological sample for this study consisted of a mixture of synthetic peptides with biologically occurring modifications combined with a yeast whole cell lysate as background, and the spectra were measured using a TripleTOF instrument. For full details on the sample preparation and acquisition see the original publication by Chalkley et al. [30]. All data was downloaded from the MassIVE data repository (accession MSV000078492).

To search the iPRG2012 data set the human HCD spectral library compiled by the National Institute of Standards and Technology (version 2016/09/12) and a TripleTOF yeast spectral library from Selevsek et al. [31] were used. First, matches to decoy proteins were removed from the yeast spectral library, after which both spectral libraries were concatenated using SpectraST [32] version 5.0 while removing duplicates by retaining only the best replicate spectrum for each individual peptide ion. Next, decoy spectra were added in a 1:1 ratio using the shuffle-and-reposition method [33], resulting in a single spectral library file containing 1 188 168 spectra.

Additionally, ANN-SoLo was used to reanalyze the human draft proteome data set by Kim et al. [34]. This large data set aims to cover the whole human proteome and consists of 30 human samples in 2212 raw files, measured using LTQ–Orbitrap Velos and LTQ–Orbitrap Elite mass spectrometers. For full details on the sample preparation and acquisition see the original publication by Kim et al. [34]. Raw files were downloaded from the PRoteomics IDEntifications (PRIDE) database [35] (project PXD000561) and converted to MGF files using msconvert [36].

To search the Kim data set the MassIVE-KB peptide spectral library (version 2018/06/15) was used. This is a repository-wide human higher-energy collisional dissociation spectral library derived from over 30 TB of human MS/MS proteomics data. The original spectral library contained 2 154 269 MS/MS spectra, from which duplicates were removed using SpectraST [32] version 5.0 by retaining only the best replicate spectrum for each individual peptide ion, resulting in a spectral library containing 2 113 413 spectra. Next, decoy spectra were added in a 1:1 ratio using the shuffle-and-reposition method [33], resulting in a final spectral library containing 4 226 826 spectra.

All MS/MS data, spectral libraries, and identification results have been deposited to the ProteomeXchange Consortium [37] via the PRIDE partner repository [35] with the data set identifier PXD013641 and via the MassIVE repository with the data set identifier RMSV000000091.4.

### 2.5 Search settings

ANN-SoLo version 0.2 was used to produce all search results. Section 3.2 compares these results to those obtained using ANN-SoLo version 0.1.3, which was previously described by Bittremieux et al. [8].

Spectrum preprocessing consisted of the removal of the precursor ion peak and noise peaks with an intensity below 1 % of the base peak intensity. If applicable, spectra were further restricted to their 50 most intense peaks. Spectra that contained fewer than 10 peaks remaining or with a mass range less than 250 *m*/*z* after peak removal were discarded. Finally, peak intensities were rank transformed to deemphasize overly dominant peaks.

The search settings for the iPRG2012 data set consist of a precursor mass tolerance of 20 ppm for the first level of the cascade search, followed by a precursor mass tolerance of 300 Da for the second level of the cascade search. The fragment mass tolerance was 0.02 Da. To evaluate the spectrum hashing performance a bin width between 0.02 Da and 1 Da and a hash length between 100 and 1600 were used. To evaluate the performance of the ANN index the number of lists was varied between 64 and 16 384 and the number of probes was varied between 1 and 1024. The number of candidates to retrieve from the ANN index was either 1024 (GPU) or 25 000 (CPU).

For the Kim data set a precursor mass tolerance of 10 ppm was used for the first level of the cascade search, followed by a precursor mass tolerance of 500 Da for the second level of the cascade search. The fragment mass tolerance was 0.05 Da. To vectorize spectra a bin width of 0.1 Da and a hash length of 800 were used. ANN searching was performed using 256 lists of which 128 were probed during searching, while retrieving 1024 candidates for each query.

All SSMs are reported at a 1 % FDR threshold.

### 2.6 Code availability

The ANN-SoLo spectral library search engine is available as a Python command-line tool. All code is released as open source under the permissive Apache 2.0 license and is available at https://github.com/bittremieux/ANN-SoLo. This web resource also includes detailed instructions on how to install and run ANN-SoLo, along with code notebooks to reproduce all analyses discussed next.

## 3 Results

### 3.1 Feature hashing converts high-resolution spectra to low-dimensional vectors

Feature hashing is used during ANN indexing to convert high-resolution MS/MS spectra to low-dimensional vectors while closely capturing their fine-grained mass resolution. Previously, 1 Da mass bins were used to vectorize the MS/MS spectra as a trade-off between fragment mass resolution and vector dimensionality [8]. However, this approach often results in multiple distinct peaks being merged into a single mass bin, leading the vector dot product to overestimate the actual spectral similarity (figure 2A, supplementary figures S1 and S2). Instead, for high-resolution spectra small mass bins should be used to closely capture the fragment masses, after which feature hashing is used to obtain low-dimensional vectors that are amenable to nearest neighbor searching (figure 2B, supplementary figures S1 and S2).

**Figure 2:**
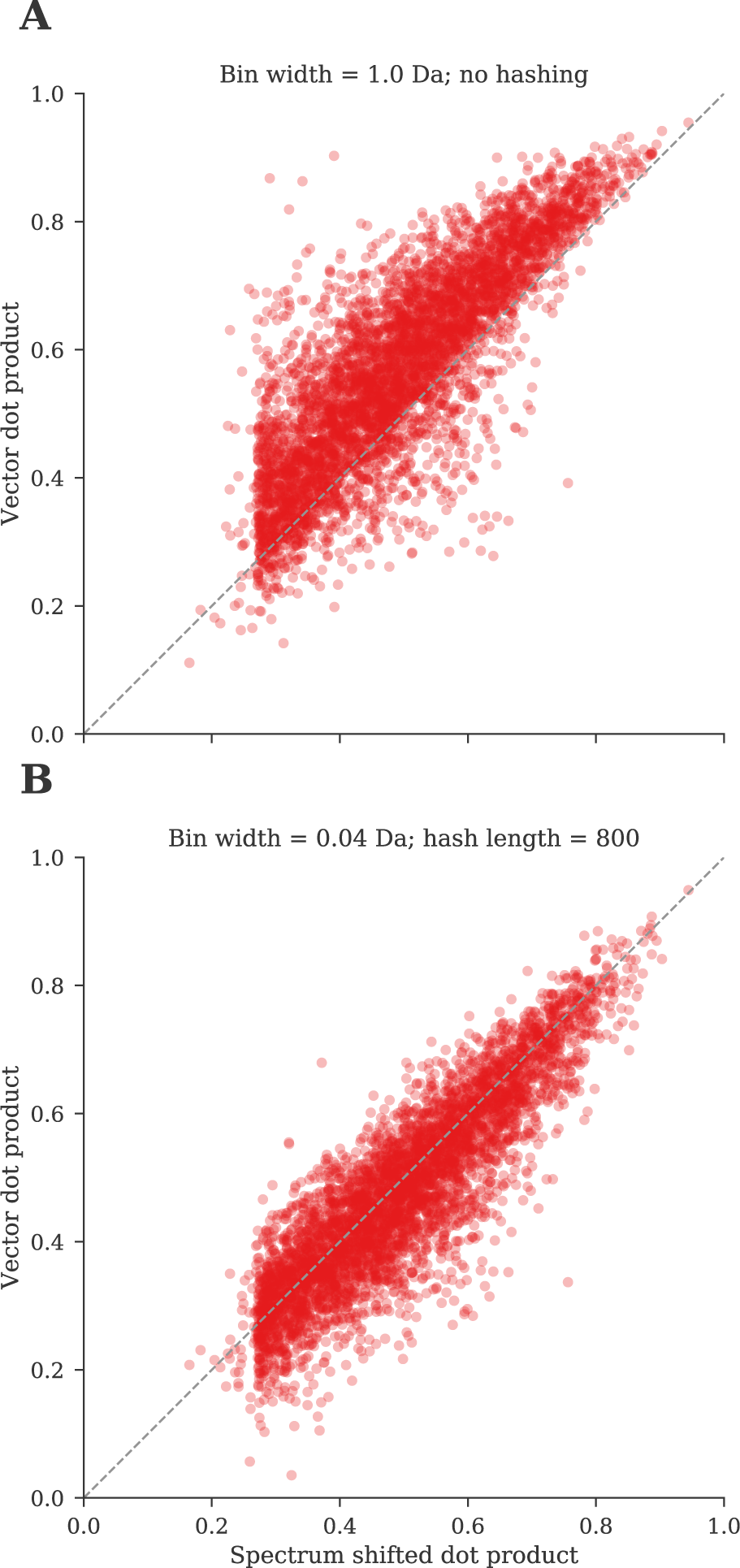
Comparison between the spectral similarity based on the spectrum shifted dot product and the vector dot product for SSMs from the iPRG2012 data set (1 % FDR). (A) The vector dot product is obtained by binning spectra using 1 Da mass bins. (B) The vector dot product is obtained by binning spectra using 0.04 Da mass bins hashed to vectors of length 800. When using 1 Da mass bins the vector dot product often overestimates the actual spectral similarity (A; SSMs above the diagonal), while small mass bins avoid spurious peak matches (B).

When feature hashing is employed, the dimensionality of the spectrum vectors is effectively disconnected from the bin width. This allows the bin width to be selected to optimally capture the fragment masses based on the operational parameters associated with the mass spectrometry run. Hashed vectors should be as short as possible to avoid the curse of dimensionality during nearest neighbor searching, while not being overly short to minimize the rate of hash collisions. Empirically, we have found that a hash length of 400 to 800 can capture the fragments in high-resolution spectra with a minimal loss of information (supplementary figures S1 and S2). Furthermore, while such hashed vectors capture the fragment resolution of high-resolution spectra more closely and enable the vector dot product to better approximate the spectral similarity, their dimensionality is actually lower than that of the original 1 Da binned vectors. Consequently, a secondary advantage of feature hashing is that these vectors require less disk space to be stored in the ANN index, and that their dot product can be computed slightly faster.

### 3.2 Highly efficient open modification searching using GPUs

To test the effectiveness of the speedups that we have introduced to ANN-SoLo, we profiled the previous and current versions of the software on the iPRG2012 data set. Although the previous version of ANN-SoLo already outperformed alternative spectral library search engines by an order of magnitude during open modification searching in terms of runtime [8], this analysis shows an additional speedup by up to an order of magnitude compared to the previously reported results (figure 3). Notably, the use of specialized GPU computing resources makes it possible to very efficiently select library candidates. The time spent during candidate selection has decreased by a factor of 30, reducing the average time required to select library candidates from 0.1222 s/query spectrum to 0.0036 s/query spectrum.

**Figure 3:**
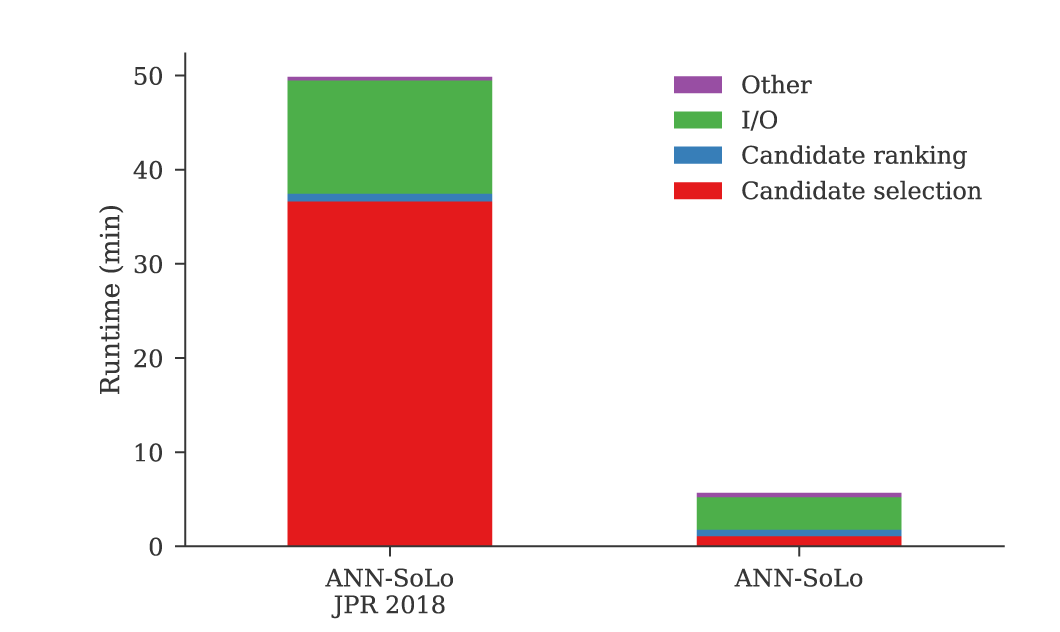
ANN-SoLo performance improvements. Whereas an open search of the iPRG2012 data set using the previous version of ANN-SoLo took 50 min, the current version performs a similar search in under 6 min. Timing results were obtained on an Intel Xeon E5-2643 v3 processor for ANN-SoLo version 0.1.3, combined with an NVIDIA GeForce RTX 2080 GPU for ANN-SoLo version 0.2.

A drawback of the approximate nature of the candidate selection step is that there is a small risk of missing the optimal library candidate. First, it is not guaranteed that the exact nearest neighbor will be retrieved from the ANN index in all cases. Two hyper-parameters (the number of lists used during index construction and the number of probed lists during querying) can be used to control the ANN searching performance (figure 4, supplementary table S1). Second, there is a discrepancy between similarity scoring in the ANN index, which is done using a standard dot product between the spectrum vectors, and the shifted dot product score matching unmodified and modified spectra to each other to obtain the final SSM ranking. Because shifted peaks are not taken into account during ANN searching, library candidates are selected based on partial matches between unshifted peaks. Consequently, if an optimal SSM contains a large proportion of shifted peaks, the corresponding library candidate will not be found via ANN searching. To alleviate this problem, multiple library candidates are retrieved from the ANN index, and these candidates are then rescored using the shifted dot product (supplementary figure S3). The number of considered library candidates is an additional hyperparameter that can be used to control the performance of ANN-SoLo by limiting the number of spectrum–spectrum comparisons that have to be performed for each query spectrum, at the expense of missing the optimal library candidate in case it is heavily modified (supplementary figure S4). A limitation of the GPU search mode is that it allows at most 1024 candidates to be retrieved from the ANN index due to GPU memory constraints. Alternatively, ANN-SoLo can be run in a CPU-only mode which does not have this limitation. This allows the user to trade off fast runtimes for a slightly higher number of identifications based on their requirements and available computational resources.

**Figure 4:**
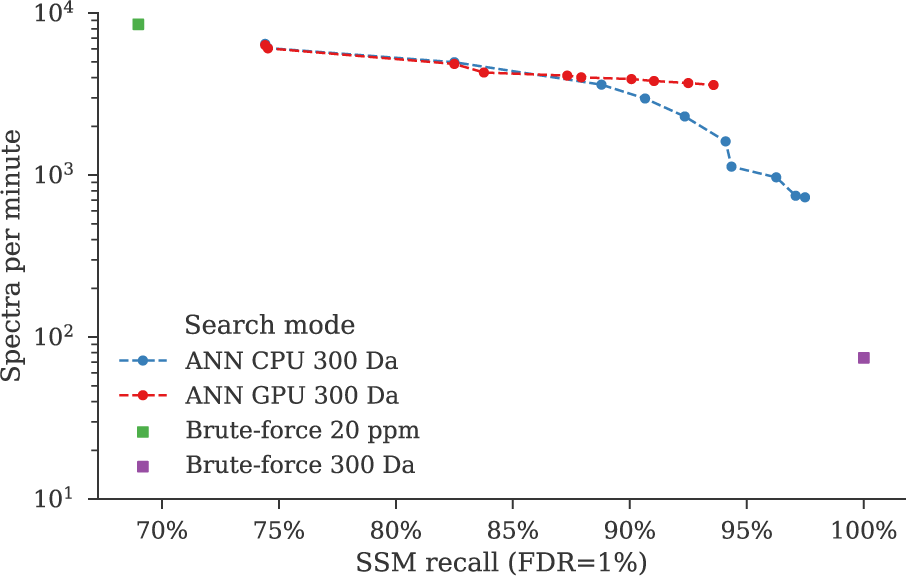
Trade-off between search speed and the number of identified spectra for the iPRG2012 data set (up and to the right is better). The number of identifications is represented as the SSM recall compared to the results of a brute-force open search without using ANN indexing. Timing results were obtained on an Intel Xeon E5-2643 v3 processor with four threads for the ANN CPU and brute-force searches, combined with an NVIDIA GeForce RTX 2080 GPU for the ANN GPU searches. Parallel execution (on the CPU or GPU) was limited to the candidate selection step. The multiple ANN results correspond to different hyperparameter configurations, with the settings that lie on the Pareto frontier shown. ANN indexing provides speed-ups of up to two orders of magnitude compared to the brute-force open search, approaching the speed of a standard search. The ANN hyperparameters can be set to achieve a higher SSM recall at the expense of a slight decrease in search speed, maximizing the number of identified spectra while still achieving a speed-up of an order of magnitude over a brute-force open search. Specific values of the ANN hyperparameters and the corresponding speed and identification performance are available in supplementary table S1.

### 3.3 Large-scale investigation of the modified human proteome

The high computational efficiency of ANN-SoLo makes it possible to perform untargeted PTM profiling via open searches at an unprecedented scale. Here we have analyzed the draft human proteome data set by Kim et al. [34], containing approximately 25 million MS/MS spectra, in combination with a large human spectral library, containing over four million spectra. A brute-force open search of such a large search space would require billions, if not trillions, of spectrum–spectrum comparisons to match all query spectra against the spectral library, which would clearly be computationally infeasible. In contrast, ANN-SoLo only needs 281 hours to search this large data set (single instance wall time), corresponding to only 8 minutes of processing time per raw file on average.

ANN-SoLo identifies over 14 million SSMs out of the 25 million query spectra. Among these identifications, approximately 9.8 million SSMs were obtained during the first level of the cascade search, and hence correspond to direct matches between query spectra and library spectra. The remaining 4.3 million SSMs have a non-zero precursor mass difference and hence represent modified peptides (figure 5 and table 1). We can see that frequently occurring modifications can mostly be attributed to various sample processing steps or can be explained by amino acid substitutions. Modifications of potential biological interest, such as acetylation, phosphorylation, GlyGly, etc., are detected at lower rates. Because these modifications are less abundant than the modifications introduced during sample processing, typically only a handful of such PTMs will be set as variable modifications during searching to minimize an unnecessary search space explosion. In contrast, our results indicate that it is possible to detect various types of biologically relevant modifications across the whole human proteome using OMS.

**Table 1:** The most frequent precursor mass differences for the Kim data set and likely modifications sourced from Unimod [38] or isotopic variants corresponding to these precursor mass differences. The delta-mass column contains the median precursor mass difference of that SSM subgroup.

| # SSMs | $\Delta m$ (Da) | Potential modification |
| --- | --- | --- |
| 9 882 777 | 0.001 |  |
| 308 387 | 57.022 | Carbamidomethyl /<br>Ala → Gln / Gly → Asn /<br>Addition of Gly |
| 246 428 | 27.996 | Formylation / Ser → Asp<br>/ Thr → Glu |
| 219 006 | 0.994 | First isotopic peak |
| 211 927 | 15.995 | Oxidation or hydrox-<br>ylation / Ala → Ser /<br>Phe → Tyr |
| 163 269 | −0.986 | Amidation |
| 133 020 | 14.016 | Methylation / Asp → Glu<br>/ Gly → Ala / Ser → Thr<br>/ Val → Leu/Ile /<br>Asn → Gln |
| 129 687 | −17.025 | Pyro-glu from Q / loss of<br>ammonia |
| 111 075 | −18.010 | Dehydration / Pyro-glu<br>from E |
| 99 286 | 1.988 | Second isotopic peak |

**Figure 5:**
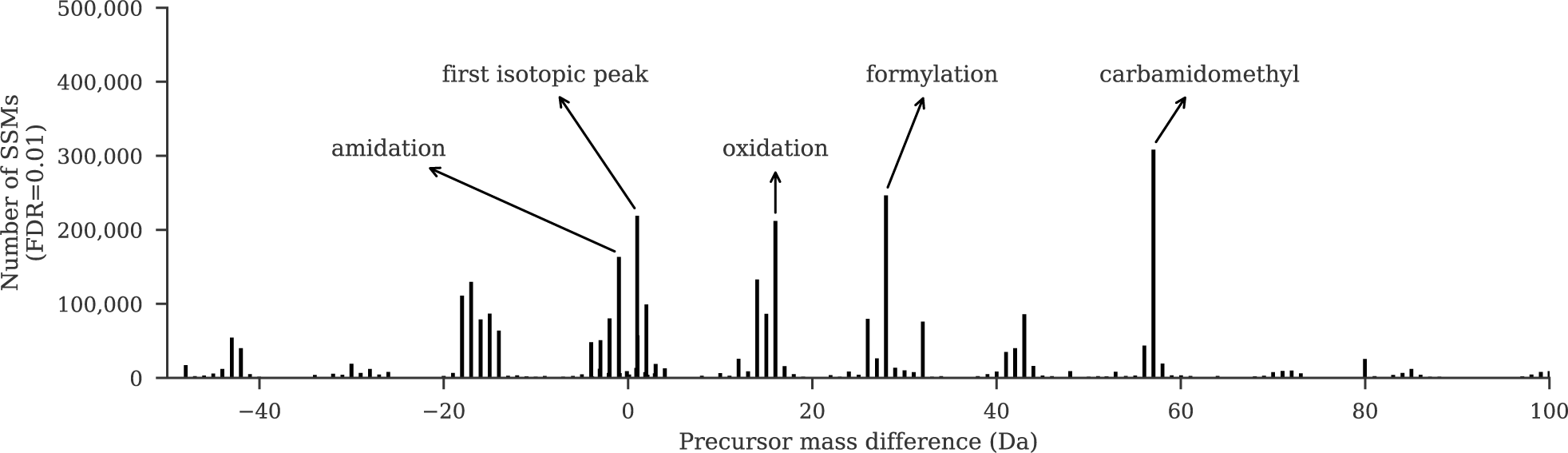
Precursor mass differences for the Kim data set (table 1). Only non-zero precursor mass differences are shown, whereas the majority of SSMs correspond to unmodified peptides with a zero precursor mass difference. The five most frequent precursor mass differences are annotated with their likely modifications.

## 4 Conclusions

We have presented an update to the ANN-SoLo spectral library search engine. ANN-SoLo uses ANN indexing to efficiently traverse the large search space encountered during OMS by selecting only a limited number of the most relevant library spectra for comparison to the unknown query spectra. We have demonstrated how specialized hardware resources, such as GPUs, can be used to optimize the candidate selection step and speed up OMS. Additionally, we have shown how feature hashing can be used to vectorize high-resolution MS/MS spectra. Feature hashing makes it possible to accurately capture the high fragment resolution of modern MS data using low-dimensional spectrum vectors.

To map mass bins in high-dimensional vectors to hash bins in low-dimensional vectors during feature hashing we have used the general-purpose MurmurHash algorithm. Alternatively, other hash functions can used as well, potentially incorporating domain knowledge. For example, a custom hash function that exploits the mass clustering effect for peptides, by not considering invalid mass values because peptides can only contain a limited number of distinct chemical elements, could be beneficial in reducing the number of hash collisions.

In this case the vectorized spectra after feature hashing were used for ANN searching to efficiently perform OMS. Via feature hashing the dimensionality of spectrum vectors can be kept low and their sparsity is reduced. As such, this technique might be used for various other downstream machine learning approaches on MS/MS spectra as well [39], because such approaches often require dense and short vectors as input.

We have demonstrated the computational efficiency and identification performance of ANN-SoLo on a large data set of the draft human proteome in combination with a repository-wide spectral library. Using traditional search engines it would be unfeasible to perform OMS on such a large volume of data. In contrast, due to its advanced, GPU-powered ANN indexing to condense the search space, ANN-SoLo can perform this task in a matter of minutes per raw file. These algorithmic advances make it possible to do OMS on a routine basis, allowing researchers to investigate the protein modification landscape at an unprecedented scale and depth.

The ANN-SoLo spectral library search engine is freely available as open source. The source code and detailed instructions can be found at https://github.com/bittremieux/ANN-SoLo.

## Acknowledgement

W.B. is a postdoctoral researcher of the Research Foundation – Flanders (FWO). W.B. was supported by a postdoctoral fellowship of the Belgian American Educational Foundation (BAEF). Some of the computational resources and services used in this work were provided by the VSC (Flemish Supercomputer Center), funded by the Research Foundation – Flanders (FWO) and the Flemish Government – department EWI. This work was supported in part by National Institutes of Health award R01 GM121818 to W.S.N.

