## Supporting information for "Extremely fast and accurate open modification spectral library searching of high-resolution mass spectra using feature hashing and graphics processing units"

### List of Tables

S1 ANN-SoLo hyperparameter evaluation . . . . . 5

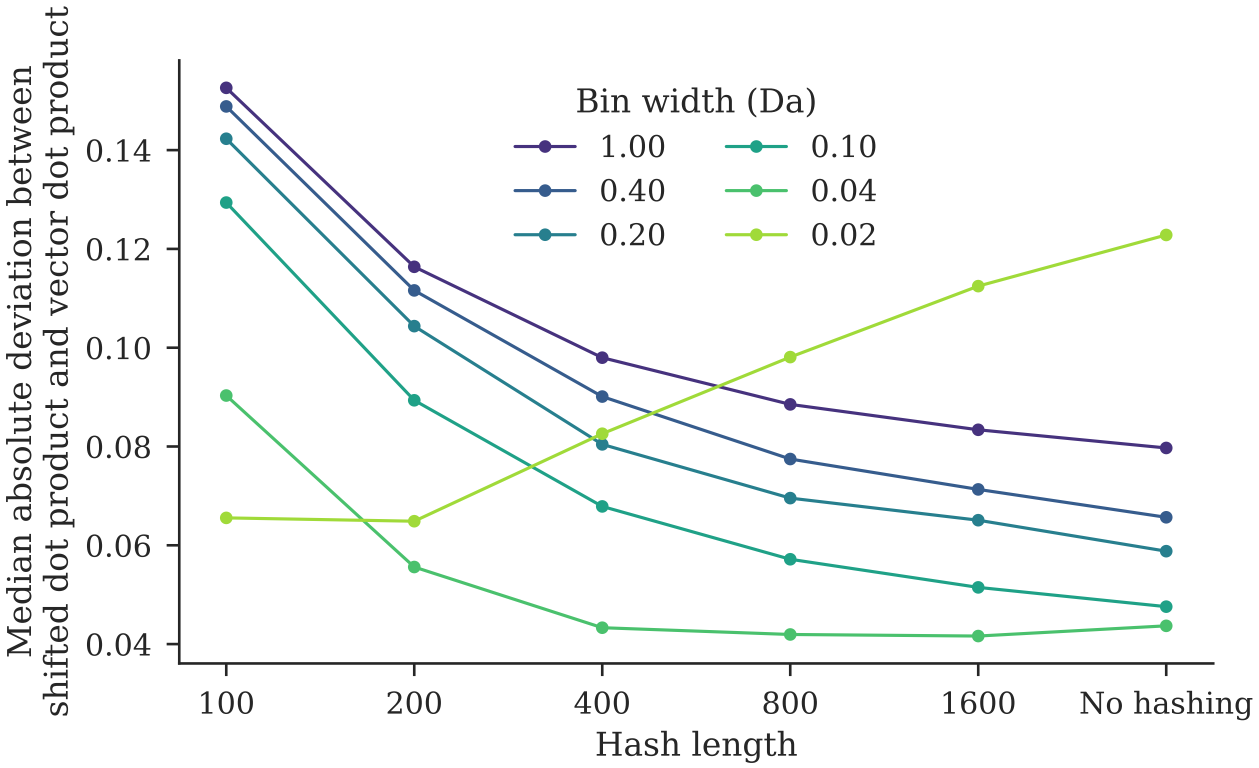

**Supplementary Figure S1:** Median absolute deviation between the peak-to-peak spectrum shifted dot product and the vector dot product. For longer hashed vectors the rate of hash collisions decreases and the vector dot product will approximate the spectrum dot product more closely. The optimal mass bin width (0.04 Da) depends on the operational parameters, with very little difference between the longer hashed vectors and the original vectors without hashing. Overly small mass bins (0.02 Da) fail to match corresponding fragments.

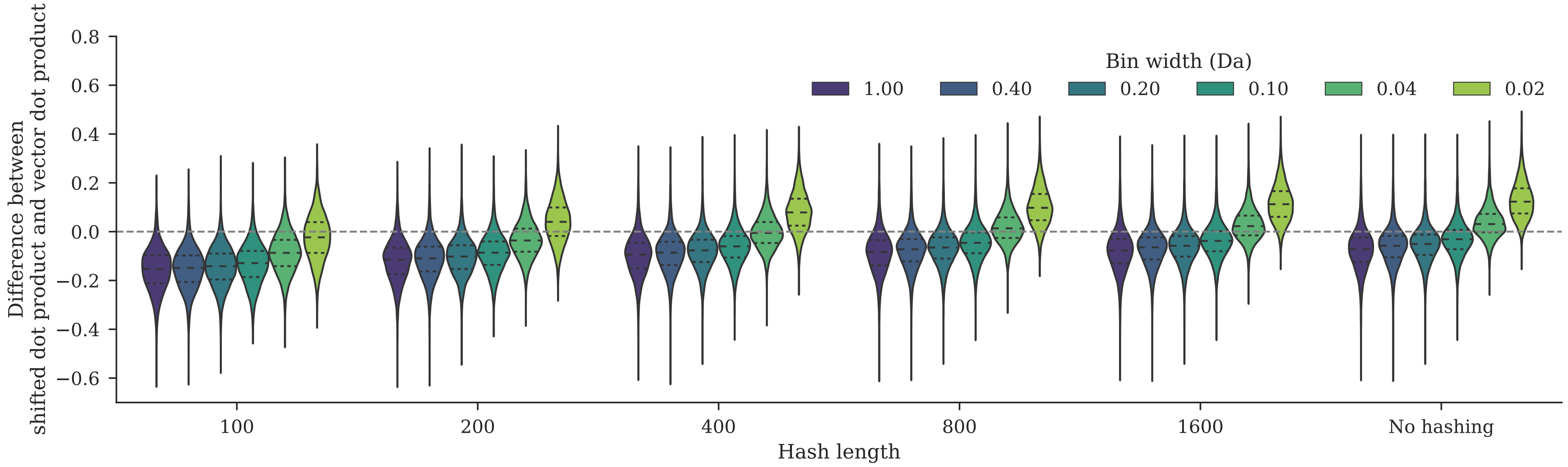

**Supplementary Figure S2:** Difference between the peak-to-peak spectrum shifted dot product and the vector dot product. Short hashed vectors suffer from more hash collisions, leading the vector dot products to overestimate the spectrum dot products (difference below 0). The spectrum shifted dot product can exceed the vector dot product (difference above 0) if the mass bins are too small (0.02 Da) or if shifted peaks due to modifications are present.

| Search mode | num_list | Build time (min) | Index size (GB) | num_probe | GPU | # SSMs | Search time (min) |
|---|---|---|---|---|---|---|---|
| Brute-force 20 ppm | | | | | | 4147 | 2.1 |
| Brute-force 300 Da | | | | | | 6011 | 241.8 |
| ANN-SoLo 300 Da | 64 | 11.2 | 3.54 | 1 | ✗ | 5024 | 8.5 |
| | | | | | ✓ | 4962 | 4.9 |
| | | | | 8 | ✗ | 5509 | 18.8 |
| | | | | | ✓ | 5382 | 4.7 |
| | | | | 32 | ✗ | 5810 | 24.3 |
| | | | | | ✓ | 5538 | 5.0 |
| ANN-SoLo 300 Da | 256 | 10.0 | 3.54 | 1 | ✗ | 4725 | 5.0 |
| | | | | | ✓ | 4714 | 4.2 |
| | | | | 8 | ✗ | 5351 | 13.0 |
| | | | | | ✓ | 5286 | 4.5 |
| | | | | 32 | ✗ | 5562 | 18.9 |
| | | | | | ✓ | 5446 | 4.7 |
| | | | | 64 | ✗ | 5784 | 21.8 |
| | | | | | ✓ | 5471 | 4.8 |
| | | | | 128 | ✗ | 5860 | 24.7 |
| | | | | | ✓ | 5625 | 5.0 |
| ANN-SoLo 300 Da | 1024 | 10.1 | 3.55 | 1 | ✗ | 4474 | 3.7 |
| | | | | | ✓ | 4471 | 3.7 |
| | | | | 8 | ✗ | 5027 | 6.4 |
| | | | | | ✓ | 5016 | 4.5 |
| | | | | 32 | ✗ | 5394 | 14.0 |
| | | | | | ✓ | 5294 | 4.6 |
| | | | | 64 | ✗ | 5501 | 17.9 |
| | | | | | ✓ | 5379 | 4.6 |
| | | | | 128 | ✗ | 5523 | 19.4 |
| | | | | | ✓ | 5387 | 4.7 |
| | | | | 256 | ✗ | 5709 | 21.6 |
| | | | | | ✓ | 5522 | 4.9 |
| | | | | 512 | ✗ | 5836 | 24.1 |
| | | | | | ✓ | 5585 | 5.1 |
| ANN-SoLo 300 Da | 4096 | 15.4 | 3.59 | 1 | ✗ | 4480 | 3.0 |
| | | | | | ✓ | 4480 | 3.0 |
| | | | | 8 | ✗ | 4956 | 4.4 |
| | | | | | ✓ | 5035 | 4.2 |
| | | | | 32 | ✗ | 5326 | 6.8 |
| | | | | | ✓ | 5285 | 4.5 |
| | | | | 64 | ✗ | 5429 | 9.6 |
| | | | | | ✓ | 5347 | 4.6 |
| | | | | 128 | ✗ | 5502 | 14.6 |
| | | | | | ✓ | 5414 | 4.6 |
| | | | | 256 | ✗ | 5538 | 18.3 |
| | | | | | ✓ | 5536 | 4.7 |
| | | | | 512 | ✗ | 5626 | 20.0 |
| | | | | | ✓ | 5472 | 4.7 |
| | | | | 1024 | ✗ | 5758 | 21.6 |
| | | | | | ✓ | 5560 | 4.9 |
| ANN-SoLo 300 Da | 16384 | 29.6 | 3.69 | 1 | ✗ | 4473 | 2.8 |
| | | | | | ✓ | 4473 | 2.8 |
| | | | | 8 | ✗ | 4959 | 3.6 |
| | | | | | ✓ | 4959 | 3.7 |
| | | | | 32 | ✗ | 5337 | 5.0 |
| | | | | | ✓ | 5249 | 4.4 |

Continued on next page

| Search mode | num_list | Build time (min) | Index size (GB) | num_probe | GPU | # SSMs | Search time (min) |
|---|---|---|---|---|---|---|---|
| | | | | 64 | ✗ | 5449 | 6.1 |
| | | | | | ✓ | 5377 | 4.6 |
| | | | | 128 | ✗ | 5551 | 7.8 |
| | | | | | ✓ | 5392 | 4.7 |
| | | | | 256 | ✗ | 5656 | 11.1 |
| | | | | | ✓ | 5544 | 4.8 |
| | | | | 512 | ✗ | 5671 | 16.0 |
| | | | | | ✓ | 5535 | 4.7 |
| | | | | 1024 | ✗ | 5786 | 18.6 |
| | | | | | ✓ | 5537 | 4.8 |

**Supplementary Table S1:** ANN-SoLo index properties and search performance for various num_list and num_probe hyperparameter combinations for the iPRG2012 data set. Timing results were obtained on an Intel Xeon E5-2643 v3 processor using four threads. Index build times include the time required to read the entire spectral library into memory and process it prior to index construction, which was around 10 minutes for the described spectral library. The reported ANN index size is the total combined size of all index files; individual files are smaller as separate files are used for different precursor charges.

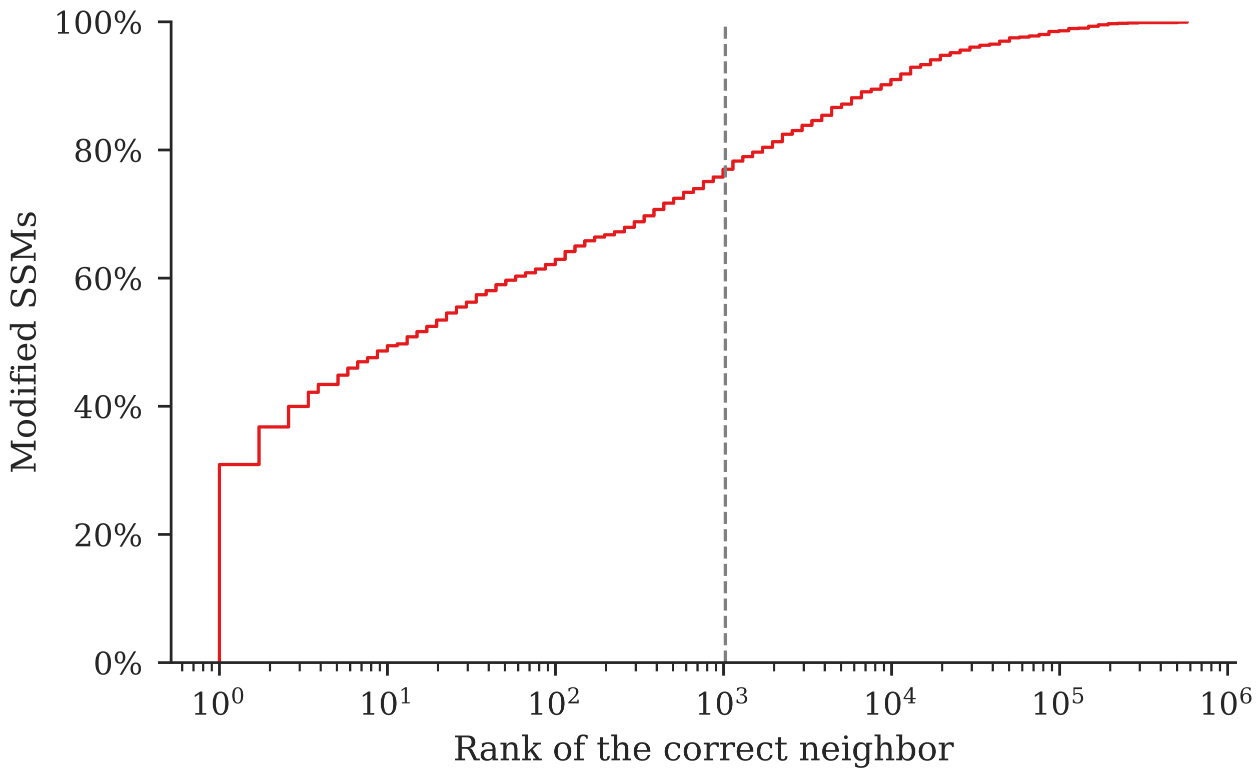

**Supplementary Figure S3:** Rank of the optimal library candidate in the ANN index for modified SSMs in the iPRG2012 data set. Library candidates are first selected in the ANN index using a standard vector dot product, without shifted peaks. Next, the SSMs are rescored using the shifted dot product to allow for modifications. However, in case the optimal SSM after rescoring requires a large proportion of peak shifting, the library candidate might have a very low rank during ANN searching, and will not be found. The dashed gray line indicates the maximum number of candidates that can be retrieved when using the GPU search mode (1024).

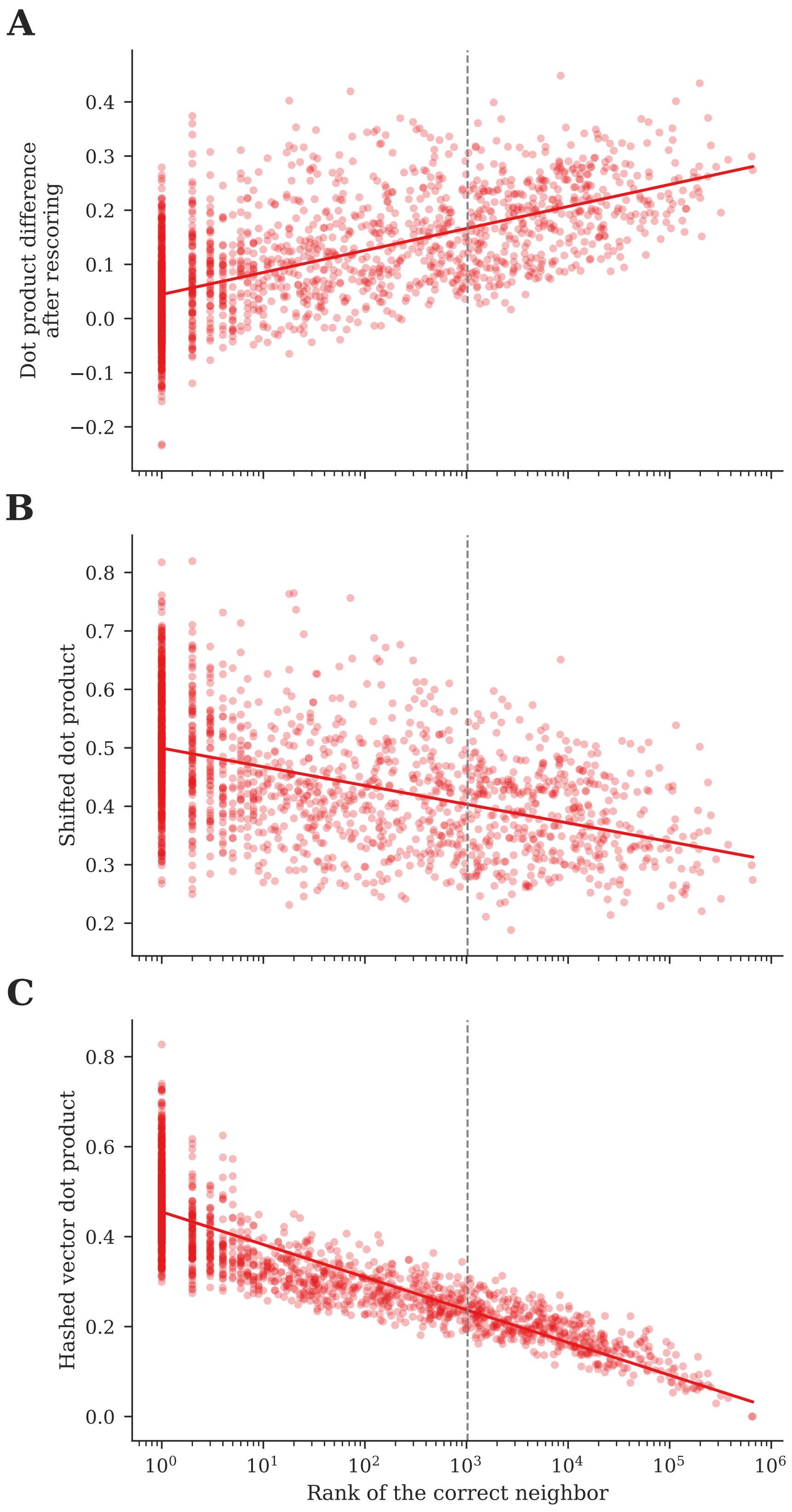

**Supplementary Figure S4:** Rank of the optimal library candidate in the ANN index versus the spectral similarity based on the vector dot product and the shifted dot product for modified SSMs in the iPRG2012 data set. (A) SSMs with a lower rank in the ANN index have a higher dot product difference after rescoring, indicating that more peak shifts take place. (B, C) SSMs with a lower rank have lower shifted dot product (B) and vector dot product (C) scores, indicating that lower-ranked SSMs that might be missed tend to be lower quality matches, irrespective of any peak shifts. The dashed gray line indicates the maximum number of candidates that can be retrieved when using the GPU search mode (1024).
